# Prognostic utility of spatial immunophenotyping in Asian breast cancer

**DOI:** 10.1101/2024.11.05.622035

**Authors:** Mohamad Hazwan Fikri Khairi, Haslina Makmur, Ju Yi Tai, Pei-Sze Ng, Chien Wen Koh, Putri Nur Fatin Amir Rudin, Iman Yi Liao, Mahmoud Khattab, Amr Ahmed, Herman Xin Yang Leong, Siti Norhidayu Hasan, Cheng-Har Yip, Pathmanathan Rajadurai, Soo-Hwang Teo, Jia-Wern Pan

## Abstract

**Background:** While tumour-infiltrating lymphocytes (TILs) are established prognostic markers in breast cancer, evidence of their prognostic value in large Asian cohorts has been limited. In this study, we evaluated the prognostic significance of immune immunohistochemistry (IHC) markers in a large Asian breast cancer dataset, including their spatial organisation and performance within multimodal machine learning models.

**Method:** A total of 664 Malaysian patients with breast cancer were included in this study. Molecular immune profiling was performed using immunohistochemistry (IHC) for CD3, CD4, CD8, and PD-L1 in 547 cases, and associations with tumour molecular features were assessed using genomic and transcriptomic data. Spatial distribution of CD8⁺ cells was subsequently characterised in a subset of 349 cases from digitised slide images using QuPath and modelled using finite mixture modelling-based pipeline (QuMix) for immunophenotype classification. To evaluate immune markers in a multivariable context, a further subset of 145 cases with complete clinical and RNA-Seq data was used to develop machine learning–based prognostic model for 10-year survival prediction.

**Findings:** Individual immune markers (CD3, CD4, CD8, and PD-L1) were not independently prognostic in the overall cohort (n=547), although CD4 and CD8 infiltration were associated with favourable outcomes in triple-negative breast cancer (n=103). Immune marker expression was linked to enrichment of immune-related pathways and lower copy number alteration burden. Notably, incorporation of immune information into machine learning models markedly improved prognostication. Whereas machine learning models incorporating traditional clinical features had an area under the receiver operating characteristic curve (AUROC) of 0.62 (95% CI: 0.37–0.62), addition of immunohistochemistry-derived and spatial immune features resulted in an AUROC of 0.71 (95% CI: 0.48–0.71), while addition of transcriptomic immune signatures achieved the highest predictive performance (AUROC = 0.88, 95% CI: 0.75–0.88). These findings demonstrate that the prognostic value of the tumour immune microenvironment in Asian breast cancer is best captured through integrated spatial and molecular immune profiling rather than by individual immune markers alone.

**Interpretation:** These results indicate that individual immune markers have limited prognostic significance in Asian breast cancer, but spatial characterization and multimodal integration improve their utility for prognostic stratification.

## Introduction

Tumour infiltrating lymphocytes (TILs) play a central role in anti-tumour immune responses in various cancers, including breast cancer, and their presence is believed to be prognostic and predictive of response to immunotherapy, chemotherapy, and other targeted therapies (de Melo Gagliato et al., 2017; El Bairi et al., 2021). In breast cancer patients, studies have shown that the density of CD8⁺ cells or TILs within the tumour region of breast tumour samples is associated with better patient outcomes, particularly in triple-negative breast cancer (TNBC) (Ali et al., 2014; de Jong et al., 2022; Liu et al., 2011; Mahmoud et al., 2011).

Other studies suggest that pre-treatment cytotoxic T lymphocytes may serve as a robust marker for predicting pathological complete response after neoadjuvant chemotherapy in breast cancer patients (Denkert et al., 2018; Hornychová et al., 2008). Moreover, increases in TILs and CD8⁺ T cell proportions in response to neoadjuvant chemotherapy are independently and more strongly associated with pathologic complete response than baseline levels of TILs (Park et al., 2020). TILs have also been heavily associated with response to immunotherapy for breast cancer patients, and various TIL-associated metrics have been proposed as biomarkers for immunotherapy, including CD8 immunohistochemistry (IHC), stromal tumour-infiltrating lymphocytes (sTILs), and immune gene expression profiles (Cocco et al., 2020; Loi et al., 2023).

Interestingly, the correlation between TILs and genomic features associated with response to immunotherapy, such as tumour mutational burden (TMB) (Samstein et al., 2019) and neoantigen load (Łuksza et al., 2017), appears to be complex. The degree and nature of clonality of somatic mutations appear to play a key role in determining whether they favor or hinder immune-mediated tumour control (McGranahan et al., 2016). In breast cancer, higher TMB has perhaps counter-intuitively been associated with lower TILs (Karn et al., 2017; Safonov et al., 2017). This is possibly the result of immunoediting, where the selection of cancer cell clones with decreased immunogenicity enables escape from immune surveillance (El Bairi et al., 2021). Other studies have also reported that neoantigen load is not correlated with CD8⁺ T-cell abundance, suggesting that TMB may be less predictive of response to checkpoint immunotherapy in breast cancer patients compared to other cancers (McGrail et al., 2021).

Importantly, recent studies have suggested that there may be differences in the response to immunotherapy for breast cancer between different populations. Subgroup analyses of the KEYNOTE-355 and KEYNOTE-522 clinical trials suggest that Asian TNBC patients may have a better response to combination immunotherapy compared to the overall TNBC population (Cortes et al., 2022; Im et al., 2024; Schmid et al., 2020; Takahashi et al., 2023). This notion is supported by genomics studies of Asian breast tumours, which suggest that Asian breast cancer patients have a more active tumour immune microenvironment relative to Western cohorts (Kan et al., 2018; Pan et al., 2024; Pan et al., 2020). However, relatively few studies of breast cancer TILs have been conducted in Asian populations, and even fewer of those have included sufficient samples to enable breast cancer subtype-specific analyses. Thus, large-scale population-specific studies of immune markers in Asian breast tumours are crucial to determine if the reported differences in tumour immune microenvironment are generalizable to the Asian population.

To date, increasing attention has been given to characterising the spatial distribution and architecture of TILs, as they have emerged as important prognostic markers in breast cancer (Cai et al., 2025; Liang et al., 2025; Walker et al., 2025). The spatial organisation of cytotoxic T cells, particularly their localisation within tumour regions such as the invasive front and tumour centre, provides additional information on tumour–immune interactions that is not captured by overall abundance alone. Spatially resolved whole-slide imaging has also enabled improved classification of immunophenotypes, including “immune-ignored”, “immune-excluded”, and “immune-inflamed” tumours. The “immune-ignored” class comprises tumours with little to no immune cell infiltration, either within the tumour or at the tumour border, while the “immune-excluded” class consists of tumours where immune cells are present primarily at the tumour border. In contrast, the “immune-inflamed” class describes tumours containing a high number of infiltrating lymphocytes (Hammerl et al., 2021). As these classifications are based on relatively subjective assessments and are not inherently quantitative, researchers have sought to standardise them by quantifying immune cell density within spatially defined tumour regions, such as the invasive front and tumour centre (Galon & Bruni, 2019; Hammerl et al., 2021; Kather et al., 2018). Nevertheless, these classifications have been associated with clinical outcomes and response to immunotherapy, underscoring their potential utility in prognostication (Hammerl et al., 2021; Wu et al., 2024).

These advances suggest that density-based assessment of individual immune markers alone may have limited prognostic value and highlight the need for more integrative approaches. In this study, we assessed the prognostic significance of IHC-based immune markers (referred to in this study as immune markers), including CD3, CD4, CD8, and PD-L1, in a large Asian breast cancer cohort and investigated their associations with relevant molecular features. We further classified samples into distinct immunophenotypes by developing a data-driven pipeline, termed QuMix, which integrates spatial characterisation of CD8⁺ cells using QuPath with finite mixture modelling. The resulting spatial features, together with immune marker density scores, were subsequently evaluated for their prognostic value within a multimodal machine learning framework. We found that IHC immune markers achieve prognostic significance in Asian breast cancer patients when evaluated within a broader multimodal and spatial framework, supporting the potential utility of these markers within clinical settings.

## Methodology

### Methodology overview

A summary of the overall study design and analysis workflow is visualized in Supplementary Figure 1. This study integrates genomic, transcriptomic, digital pathology, and clinical data to evaluate the prognostic relevance of immune markers in a large Asian breast cancer cohort. Formalin-fixed, paraffin-embedded (FFPE) tumour samples were subjected to immunohistochemical staining for CD3, CD4, CD8, and PD-L1, followed by semi-automated quantification. Prognostic significance of these marker scores was assessed using adjusted Cox proportional hazards models, while associations with tumour molecular features and related pathways were tested using univariable and multivariable regression.

Beyond IHC scores, spatial characterization of CD8⁺ T-cell infiltration was performed using QuPath to derive spatial immune features. A data-driven framework, QuMix, was developed using finite mixture modelling to stratify samples into immune-inflamed, immune-excluded, and immune-ignored immunophenotypes, and subsequently validated against a manual scoring approach. Finally, to evaluate these markers within a multivariable framework, they were integrated into a previously developed ensemble machine learning model to classify patients according to 10-year prognosis.

### Study cohort and sample selection

Our study cohort consists of 547 Malaysian women with breast cancer recruited from the Subang Jaya Medical Center between 2012 and 2018 as part of the MyBrCa Genetics study (Tan et al., 2018). For this study, we conducted a retrospective immunohistochemical analysis of FFPE breast primary tumour tissue collected during surgery. These analyses were paired with overall survival, clinical, and transcriptomic data collected from fresh frozen tumour samples as part of the MyBrCa study that have been described previously (Pan et al., 2020). Patient enrolment and analyses of patient samples were approved by the Ethics Committee of Subang Jaya Medical Center (reference no: 201109.4 and 201208.1) and written informed consent was provided by each patient.

### FFPE tissue sectioning and staining

Immunohistochemical (IHC) stained FFPE tissue sections were prepared as per a standard diagnostic workflow. Four slides per tumour sample were stained with primary antibodies targeting specific antigens: anti-CD3 (clone 2GV6, predilute; Ventana Medical Systems), anti-CD4 (clone SP35, predilute; Ventana Medical Systems), anti-CD8 (clone SP57, predilute; Ventana Medical Systems), and anti PD-L1 (clone SP263, predilute; Ventana Medical Systems), with an additional slide stained for with hematoxylin and eosin (H&E). The antibody staining was conducted using a Ventana Bench Mark XT Autostainer, and stained slides were digitised using an Aperio AT2 whole slide scanner.

### Genomic and transcriptomic data

Detailed descriptions of the genomic and transcriptomic features of this cohort have been described in our previous studies (Pan et al., 2024; Pan et al., 2020). In brief, the features were derived from paired-end short-read Illumina whole-exome, whole-transcriptome, and shallow-whole genome sequencing of tumour and blood-derived matched normal samples from each patient, mapped to the hs37d5 reference genome. The features analysed in this study include somatic variants, single-base-substitution (SBS) mutational signatures, and homologous recombination deficiency (HRD) scores derived from the whole-exome data, copy number aberrations (CNAs) derived from shallow whole-genome data, and gene set-based pathway expression scores derived from whole-transcriptome data.

### Semi-automated scoring of immunohistochemistry-based immune markers

For CD3, CD4, and CD8 scoring, Aperio Imagescope was used to view the scanned images and delineate 50 regions of interest (ROI) using annotation boxes, each of size 0.7 mm x 0.4 mm, within the borders of the invasive tumour region. The borders of the invasive tumour region were determined with reference to the H&E-stained slides. Then, the Pixel Positive v9 algorithm implemented in Aperio Imagescope was used to calculate the number of positively stained pixels within the annotation boxes at a 0.16 colour saturation threshold. The number of positive pixels was then divided by the total number of pixels within the annotation boxes to obtain the percentage of the annotated area that was positively stained for CD3, CD4 and CD8. For PD-L1 scoring, we used the combined positive score (CPS) system to score each sample as a 1 (positive) or 0 (negative) for PD-L1 expression based on CPS>1.

### Pathway analysis

Genes belonging to the biological pathways included in our analysis were obtained from the Kyoto Encyclopedia of Genes and Genomes (KEGG) database (Kanehisa & Goto, 2000). The following KEGG pathways were analysed: hsa04623 (cGas-STING), hsa04350 (TGF-beta signalling), hsa04660 (T-cell receptor signalling) and hsa04612 (antigen processing and presentation). Gene Set Variation Analysis (GSVA) of each pathway was performed using the GSVA package (v.1.48.3) implemented in R on the pathway gene sets, with gene expression data from the MyBrCa cohort used as the input data (Hänzelmann et al., 2013). The gene expression data was normalized prior to input into GSVA using “trimmed mean of M-values” (TMM) and “variance modeling at the observational level” (voom) normalization as implemented in the edgeR (v. 3.20.9) and limma (v. 3.34.9) packages in R (Chen et al., 2025; Ritchie et al., 2015).

### Spatial characterization of CD8⁺ cells using QuPath

#### Cell segmentation

The detailed protocol for semi-automated quantitative scoring using QuPath (v. 0.3.2) has been described previously by Berben et al. (2020) (Supplementary Figure 2). The tumour border for each sample was manually annotated using the *Polygon* tool, which was used to delineate the tumour and normal regions from a whole-slide image. A script provided by the QuPath developer (*QuPath-Creating tumor margin annotations.groovy*) was then used to demarcate the annotated tumour regions into three sections: the outer margin, inner margin, and tumour center (TC) (Bankhead, 2018).The tumour invasive front (IF) was defined as the combined area of the inner margin (500 µm wide) and the outer margin (500 µm wide). Next, the cells within the IF and TC were segmented using the QuPath *Cell Detection* tool. The settings used for cell detection are summarized in Supplementary Table 1. Following detection, cell segmentation was refined with the QuPath “*smooth object features”* function using default settings.

#### Cell classification

For cell classification, a classifier was trained to classify the segmented cells using control tissues stained with the same markers that were included with each whole-slide image in our cohort using QuPath’s *Train Object Classifier* tool. For each whole-slide image, at least three different manually annotated regions from the control staining were used to train the QuPath model for cell classification. In order to train the classifier, a rectangle was drawn over the control region and the cells within this region were segmented using the QuPath *Cell Detection* tool (Supplementary Figure 2B). Then, within this annotated region, a group of stained segmented cells was manually outlined using the *Polygon* tool and labelled as “Immune cells”, while the unstained cells were labelled as “Others”, and used to train the classification model. The model was then applied to segmented cells in the tumour region, classifying some stained cells as “Immune cells” and all remaining cells as “Others”. If training on the control region precluded accurate classification of cells within the tumour region (based on visual inspection), then a classifier trained on a different sample was used, and if that was still not accurate, then the classifier was trained using the tumour region itself. The settings used with the *Train Object Classifier* tool during classifier training were the default settings except for the *measurement features* setting where only *smoothed featur*es were selected. Cell classification was further refined using the QuPath *Cell Intensity Classification* tool. The tool was used further differentiate between positively- and negatively-stained immune cells by selecting the *smoothed features* measurement with the options “Smoothed: 25 µm: Nucleus: DAB OD mean” and the resulting automatic threshold was further adjusted manually. The final numbers of classified immune cells (positive and negative) and other types of cells were then exported out.

### Immunophenotype classification using finite mixture model–based thresholding and manual validation

The QuMix pipeline was developed to classify immunophenotypes using spatial immune features derived from QuPath and data-driven thresholding based on finite mixture modelling (FMM) (Supplementary Figure 2F). FMM was chosen as it is well suited for spatial immune features, which often exhibit continuous and overlapping distributions without clear empirical thresholds. FMM enables data-driven identification of latent subpopulations, reducing reliance on arbitrary cut-offs and improving the reproducibility of the classification. The features used for classification here included the average CD8⁺ cell density in the invasive front (IF) and tumour center (TC), as well as the corresponding IF-to-TC density ratio. In this study, the IF-to-TC density ratio was implemented to better distinguish tumours with immune cells restricted to the IF (immune-excluded) from those with infiltration into the TC (immune-inflamed), which cannot be resolved solely on absolute density. The FMM analysis was performed using the “normalmixEM” function from the “mixtools” package in R (v2.0.0) (Benaglia et al., 2009) using k=2 on a single dimension. Thresholds were derived separately for each feature (CD8 TC, CD8 IF, and IF/TC ratio) with the cut-off defined as one standard deviation above the mean of the lower component distribution. Samples were first classified as immune-ignored if the average CD8⁺ cell density in both TC and IF was below the defined density threshold. The remaining samples above this threshold were further stratified based on the density ratio, where values above the ratio threshold were classified as immune-excluded, and those below were classified as immune-inflamed.

QuMix classification was validated against immunophenotype classification based on a manual scoring method previously described in Hammerl et al. (2021), referred to in this study as the Hammerl method. In brief, this method quantifies CD8⁺ cells by calculating their percentage in the tumour border and center regions. Similar to QuMix, the percentage mean at the border and center, as well as the ratio of the percentage mean at the border to that at the center, were used as input features for finite mixture modelling to derive classification thresholds.

### Machine learning–based prognostic modelling using digital pathology-derived immune features

The development of the machine learning framework used in this study has been previously described in Khairi et al. (2025). The dataset in that study was derived from the MyBrCa cohort and included patients with available clinical information (tumour grade, age at diagnosis, tumour size, number of positive lymph nodes, and HER2/PR/ER receptor status), survival data, and RNA-seq data. Patients with stage 4 disease were excluded from this cohort. Detailed preprocessing of the RNA-seq data has also been previously described in Khairi et al. (2025). The dataset was then labelled using a 10-year survival threshold. Specifically, patients who survived more than 10 years after diagnosis were considered to have a good prognosis, whereas those who succumbed to the disease within 10 years were considered to have a poor prognosis. Accordingly, patients who were still alive but had less than 10 years of follow-up were excluded from the dataset. This classification enables a binary modelling framework while providing a clinically interpretable endpoint and reducing bias from right-censoring among patients with shorter follow-up.

The machine learning framework was developed in R (v4.3.3) using the machine learning framework available in the “caret” (v7.0-1) package (Kuhn et al., 2008). The input features are summarized in Supplementary Table 2, where digital pathology-derived immune features included QuPath-derived CD8⁺ T-cell densities and immunohistochemistry-based immune marker scores and PD-L1 expression, clinical features included key clinicopathological variables, and gene expression features comprised of a 77-gene signature derived from the MyBrCa cohort (previously described in Khairi et al. (2025)). The machine learning architecture consists of an inner and outer loop for dataset splitting and training–testing, using a pipeline based on Random Forest (RF) and Support Vector Machine (SVM) algorithms to predict the probability of a patient having a good prognosis. (Supplementary Figure 3) Three SVM kernels: linear, radial, and polynomial kernels, were implemented in this framework. The predicted probabilities from these four models (one RF and three SVMs) were averaged to generate a single ensemble prediction for each sample. The dataset splitting was performed in the outer loop using a one-fold random stratified split under a 7:3 ratio, which generated five distinct training and testing datasets. Prior to model training, each training dataset undergoes z-score normalization and collinearity removal within the inner loop.

In every iteration of model training, a five-fold cross-validation was applied for hyperparameter tuning, using default tuning grids to identify the optimal parameters that maximized the area under the receiver operating characteristic curve (AUROC). Overall, a total of 20 sub-models were trained (5 iterations × 4 models per iteration). Model performance was evaluated using AUROC on the corresponding hold-out test sets, and the final model was selected based on the median AUROC across all iterations.

### Survival and statistical analysis

Overall survival data were obtained from the Malaysian National Registry by matching unique national identity card numbers. Overall survival was defined as the time from diagnosis to death for deceased patients, or to the last registry update for patients without a recorded death, who were assumed to be alive at last follow-up. Survival analyses were performed using Cox proportional hazards models using the “coxph” function from the survival package in R (v3.5-5) (Borgan, 2001). Adjusted survival curves were generated using the “ggadjustedcurves” function with the conditional approach from the “survminer” package (v0.4.9) (Kassambara et al., 2016).

Associations between immune markers and molecular features were evaluated using univariable and multivariable regression analyses. Univariateble regression was done using the generalized linear models (GLM) with a Gaussian distribution, implemented using the “glm” function in the R base stats package to assess the relationship between individual immune markers (CD3/CD4/CD8) and the tumour molecular variables. Multivariable analyses were conducted using standard linear regression models implemented via the “lm” function to incorporate all of the tumour molecular variables simultaneously. For variables that had more than a 0.8 correlation with other variables, only the variable with the highest correlation with the immune marker being investigated was selected to be included in the multiple regression analysis.

Chi-square tests were conducted to compare categorical variables between groups, while Student’s t-tests were used to compare continuous variables between groups. Associations between two continuous variables were examined using Pearson’s correlation. P-values of 0.05 was used as the cutoff for statistical significance. Confidence intervals (CI) reported are for the 95% confidence threshold.

## Results

### Study populations and clinicopathological characteristics

The study cohort consisted of 664 female patients with breast cancer recruited sequentially at the Subang Jaya Medical Center (SJMC) who were part of the Malaysian Breast Cancer (MyBrCa) Genetics study. The mean age at diagnosis was 54.0 ± 11.9, with most of cases having histologically confirmed invasive ductal carcinoma (88.4%). Most tumours were either grade 2 (42.2%) or grade 3 (48.1%)), and either stage II (46.5%) or stage III (29.9%). By immunohistochemistry, 287 (49.8%) were classified as HR+/HER2-, 63 (10.9%) were classified as HR+/HER2+, 88 (15.3%) were classified as HR-/HER2+, and 103 (17.9%) classified as triple-negative breast cancer (TNBC) (Supplementary Table 3).

For downstream analyses, the cohort was further subset based on data availability. CD3, CD4, and CD8 immune marker scores were quantified in 547 cases using the semi-automated Aperio ImageScope platform, while spatial immune profiling using the QuPath-based QuMix pipeline was performed in 349 cases. On the other hand, the cohort used to develop the machine learning-based prognostic model comprised of 145 patients and was derived from the overlapping subset of cases with both IHC and QuMix data that also met additional inclusion criteria described previously in the Methodology section (Supplementary Table 3).

### Clinicopathological correlates and prognostic significance of immune markers in Asian breast cancer

For each of the 547 breast cancer patients, serial FFPE tissue sections of primary tumour samples were stained by immunohistochemistry for CD3, CD4, CD8 to obtain digitized whole-slide images and scored using a semi-automated method to obtain the percentage of the tumour region that was stained with the marker for each sample. PD-L1 was similarly stained and scored using the combined positive score (CPS) system with a CPS>1 cutoff. The raw scores for CD3, CD4 and CD8 were then log-transformed into a normal distribution (Supplementary Figure 4) for all subsequent analyses. Next, we divided our patients into high- and low-scoring groups for each immune marker according to their median values to compare their clinicopathological characteristics, which are summarized in the Supplementary Table 4. Patients with higher CD3 or CD4 scores were more likely to have higher tumour grade (P = 6.1e-6 and 0.0013, respectively), have tumours that lack the estrogen receptor (ER) and progesterone receptor (PR), and have HER2-positive tumours. Patients with higher CD8 scores were more likely to be younger, but there was no association with immunological staining of ER, PR or HER2. Overall, PDL1 positivity was low (82 out of 547 (15.0%) scored CPS>1), and where positive, it was associated with younger age (P = 0.0127), higher grade and estrogen- and progesterone-receptor negativity. The TILs were not associated with tumour stage, tumour size, nor histological subtype.

The median follow-up time for patients was 67.7 months, with 114 out of 547 patients (20.8%) having passed away during follow-up. Applying a Cox proportional hazard model to the patient subset (n=574), adjusting for tumour grade, tumour stage, and clinical subtype, we found clear differences in overall survival between clinical subtypes, with HR+/HER2+ patients having the best overall survival, and TNBC patients having the worst overall survival (Supplementary Figure 5a). We also found significant differences in overall survival when comparing the different cancer stages, with each increase in stage being associated with a hazard ratio of 2.64 (CI: 2.01-3.47) (Supplementary Figure 5b-c).

We then compared the immune IHC scores to the overall survival data for our patient cohort, adjusting for tumour stage, tumour grade, HR status, and HER2 status using Cox proportional hazard models. PD-L1 positivity was not associated with any difference in overall survival (P = 0.85; Figure 1). This lack of association remained consistent across subgroup analyses stratified by receptor status, including triple-negative breast cancer, hormone receptor–positive (HR+), and HER2+ (HR−/HER2 +) subtypes. Similarly, higher CD3, CD4, and CD8 scores were not significantly associated with overall survival, although trends towards improved survival were observed for CD4 and CD8 (Figure 1). In patients with triple negative breast cancer (n = 103), after adjusting for tumour grade, tumour stage and chemotherapy treatment, we found that high CD4, and CD8 scores were independently associated with better overall survival, with larger differences in overall survival between the high- and low-scoring groups at five years compared to the analysis using the full cohort (Figure 1). In patients with HR+ subtypes (n = 350), after adjusting for tumour grade, tumour stage and hormonal treatment, we found that only high CD3 scores was positively association with overall survival (P = 0.030; Figure 1). In HER2+ patients (n = 151), after adjusting for tumour grade, tumour stage and hormonal therapy (we were unable to adjust for HER2-targeted therapy due to incomplete data), we found no significant associations between the immune markers and overall survival (Figure 1).

**Figure 1.**
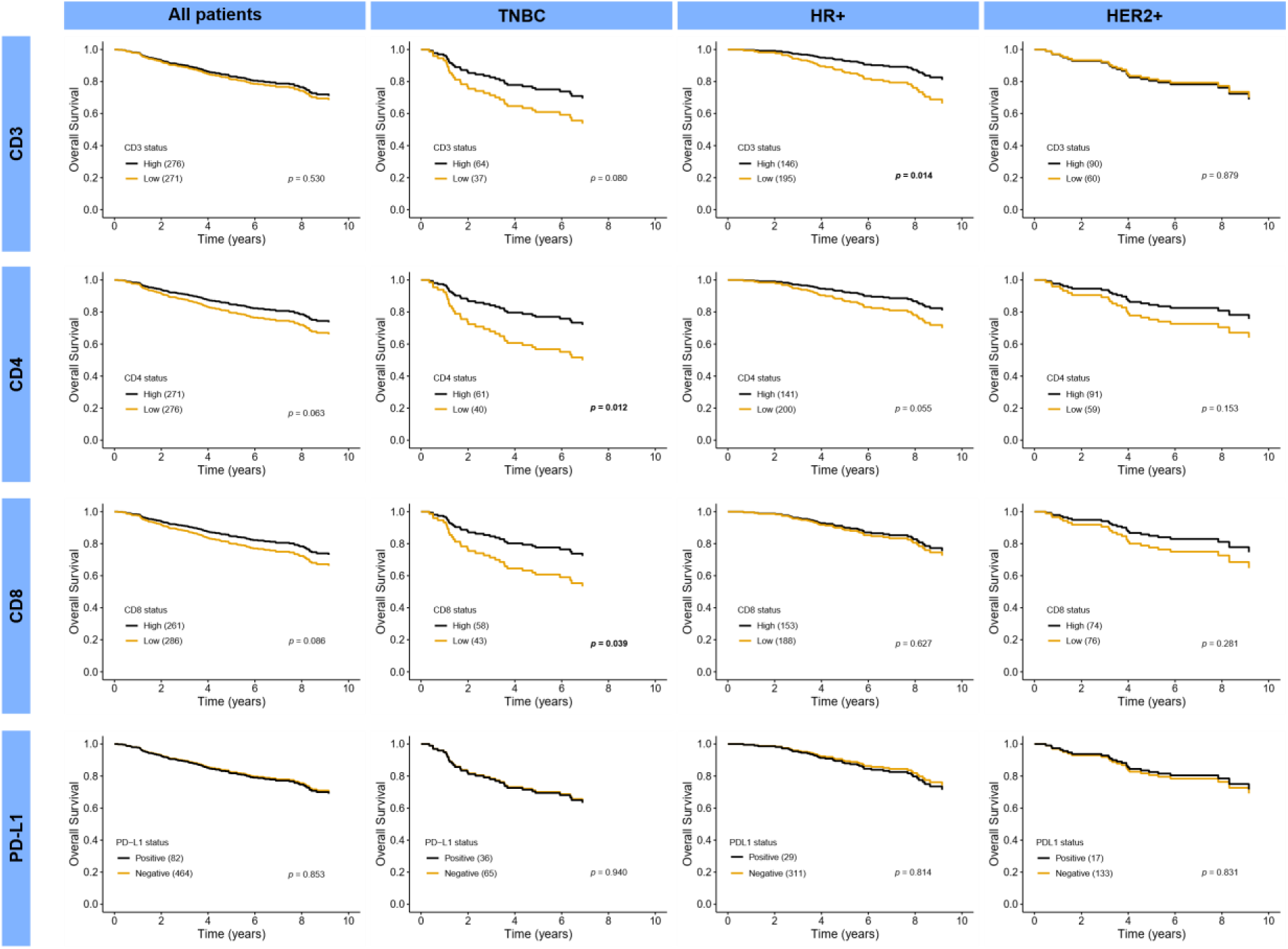
Overall survival by immune marker status in breast cancer subgroups. Overall survival of all breast cancer patients, TNBC patients, HR+ patients, and HER2+ patients stratified by high versus low CD3, CD4, or CD8 scores (based on median value), or positive versus negative PD-L1 based on CPS>1. Sample sizes for each group are indicated in brackets. Overall survival was adjusted using a Cox proportional hazard model that also included tumour stage, tumour grade, HR status, and HER2 status as variables. P-values shown are the p-values of the variable from the Cox proportional hazard model.

### Correlation of genomic and transcriptomic features with the immune markers

Next, we compared the immune IHC scores to genomic and transcriptomic features of the cohort derived from whole-exome, whole-transcriptome, and shallow whole-genome sequencing. We found that high CD3 scores were associated with TP53 somatic mutations, high APOBEC-associated single-base-substitution (SBS) signatures 2 and 13, DNA repair–associated SBS signatures 3 and 8, while negatively associated with clock-like SBS signatures 1 and 5 (Figure 2a). On the other hand, high CD4 scores were associated with high HRD scores and high APOBEC-associated SBS signatures 2 and 13. All 3 immune markers were also negatively associated with CNAs (Figure 2a).

**Figure 2.**
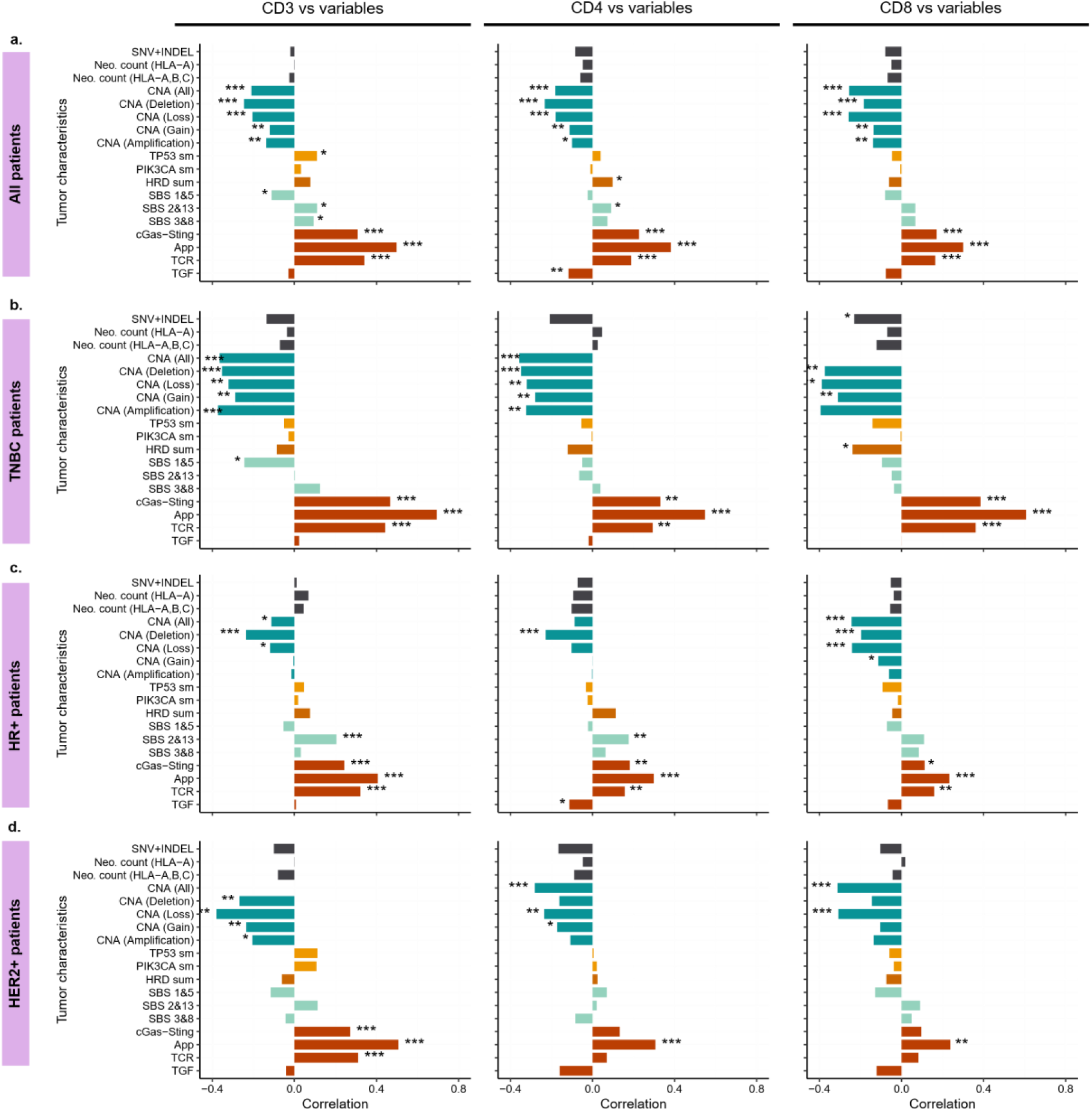
Correlation of CD3, CD4, and CD8 staining with tumour molecular features. The tumour molecular features include SNV and indel counts (SNV+Indel), HLA-A only or HLA-A, -B, and -C neoantigen counts, copy number aberrations (CNA) including copy number deletion, loss, gain, and amplification, TP53 and PIK3CA somatic mutations (sm), HRD sum scores from scarHRD, COSMIC mutational signatures for aging/clock-like (SBS 1&5) and APOBEC (SBS 2&13), and gene set variation analysis (GSVA) scores for four KEGG pathways – cGAS-STING signalling (cGAS-STING), antigen processing and presentation (APP), T-cell receptor signalling (TCR), and TGF-β signalling (TGF). Comparisons were conducted for all patients (a.) as well as for TNBC patients only (b.), HR+ patients only, (c.) or HER2+ patients only (d.). Correlation shown is Spearman rank’s correlation. Asterisks indicate significant correlations (* indicates p < 0.05, ** indicates p < 0.01, *** indicates p < 0.001).

From the transcriptomic data, we derived gene set variation analysis (GSVA) scores for four KEGG pathways that have known associations with cancer immunity: cGAS-STING signaling (cGAS-STING), antigen processing and presentation (APP), T-cell receptor signaling (TCR), and TGF-β signaling (TGF-β). All three immune markers had strong positive correlations with the cGAS-STING signaling, antigen processing and presentation, and T-cell receptor signaling pathways (Figure 2a). We also repeated the analysis in the TNBC, HR+, and HER2+ subgroups of patients to determine if there were any unique subgroup-specific associations between immune markers and genomic or transcriptomic features. For the most part, the trends seen in the overall analysis were also found in the subgroup analyses, but there were some important differences. In the TNBC subgroup analysis, the negative correlation between immune markers and CNAs was stronger relative to the overall analysis (i.e. rs = -0.35 in TNBC subgroup vs rs = -0.18 overall for all CNAs vs CD3), and HRD scores had a stronger negative correlation with CD8 scores (Figure 2b). On the other hand, the positive correlation between immune markers and the cGAS-STING, APP, and TCR pathways is also stronger relative to the overall analysis (Figure 2b). In the HR+ subgroup, the negative association between immune markers and CNAs is weaker relative to the overall analysis, but the correlation with the APOBEC-associated SBS 2 and 13 mutational signatures was stronger for CD3 and CD4 (Figure 2c). For the HER2+ subgroup, the negative correlation between immune markers and CNAs and the positive correlation between immune markers and the cGAS-STING, APP, and TCR pathways were present, but there were no significant correlations between immune markers and any other genomic features (Figure 2d).

We further validated these findings by conducting univariable as well as multiple linear regression analyses for each of the immune markers, with all the genomic and transcriptomic features listed above included as independent variables. For the multiple regression analyses, autocorrelated features (features with a greater than 0.8 correlation with each other) were manipulated such that only the feature with the higher correlation with the dependent immune marker variable was included. Our regression analyses were largely in agreement with our previous results, as we found that SNVs, indels, and predicted neoantigen load had no association or negative associations with the immune markers (Supplementary Table 5). Indeed, for our multiple regression analyses, only tumour subtype and expression of the APP, TCR, and TGF-β pathways were predictive of CD3 scores, only expression of the APP pathway was predictive of CD4 scores, and only tumour subtype and CNAs were predictive of CD8 scores.

### CD8-based digital classification of tumour immunophenotypes using the semi-automated QuMix pipeline

We then assessed the utility of CD8⁺ immunohistochemistry for classifying tumours into distinct immunophenotypes in our cohort by developing a semi-automated pipeline that combines an open-source biomedical image analysis tool with finite mixture model analysis, referred to in this study as QuMix. QuMix classifies tumours into immune-ignored, immune-excluded, and immune-inflamed categories using the readily available digital pathology tool QuPath combined with mixture model analysis. Classification was performed on 349 CD8-stained Finite mixture modelling of CD8⁺ T-cell density at the TC and IC as well as the IF-to-TC density ratio, suggested a predominantly unimodal distribution, which was then used to derive thresholds for immunophenotype classification (Figure 3b).

**Figure 3.**
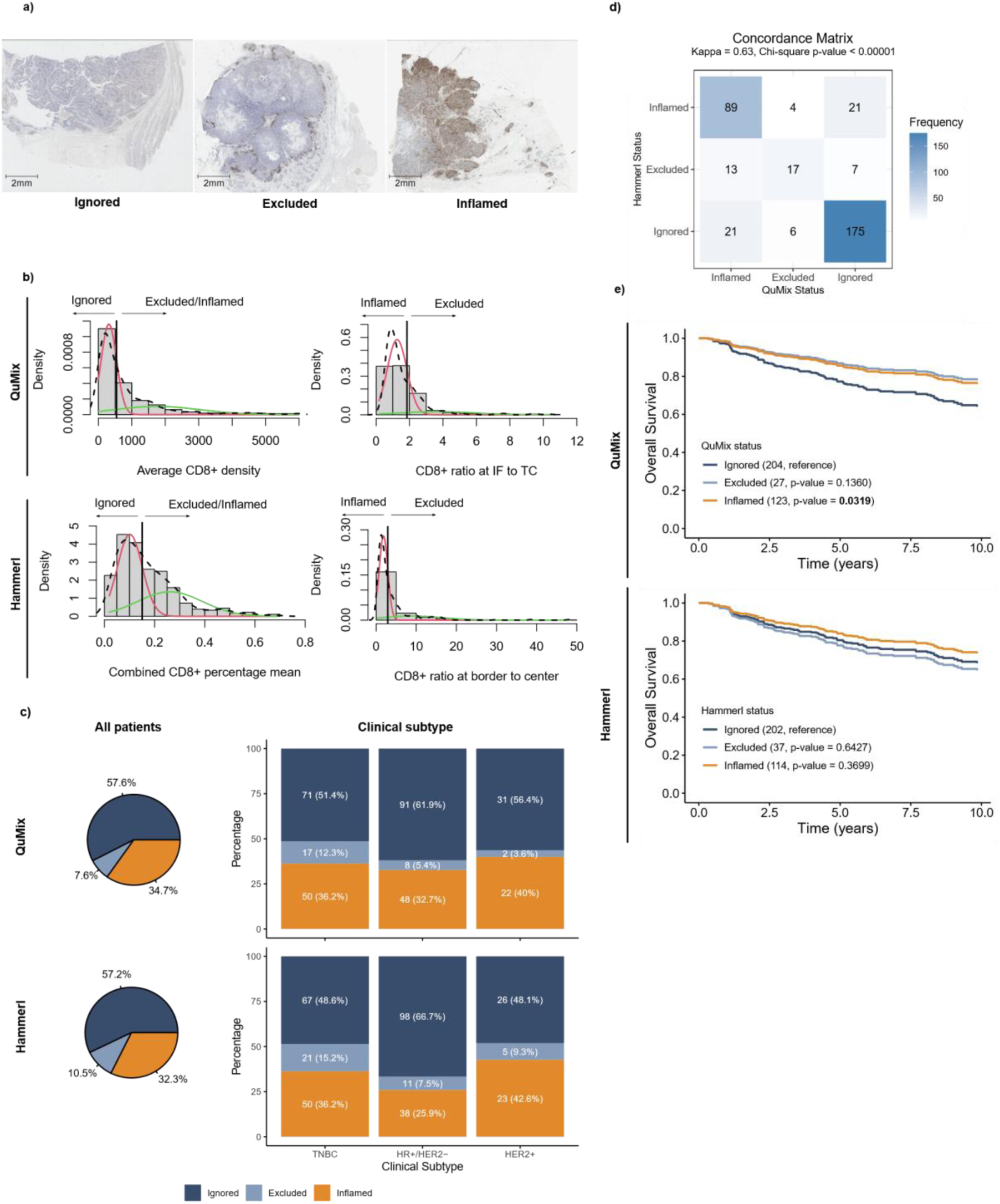
Spatial immunophenotype classification and comparison between QuMix and Hammerl approaches. a) Representative tissue images illustrating the three immunophenotype categories. b) Finite mixture modelling used to determine classification thresholds for both QuMix and Hammerl methods. c) Proportion of patients assigned to each immunophenotype category by QuMix and Hammerl across all patients and within molecular subtypes (TNBC, HR+/HER2−, HER2+). d) Concordance matrix comparing immunophenotype classifications between QuMix and Hammerl methods. e) Survival curves based on immunophenotype classification, adjusted using a Cox proportional hazards model including hormone receptor (HR) status, HER2 status, tumour grade, tumour stage, and tumour size as covariates.

Based on thresholds derived from mixture model analysis (Figure 3c), 57.6% of patients were classified as immune-ignored, followed by immune-inflamed (34.7%) and immune-excluded (7.6%) (Figure 3c). A similar distribution was observed across molecular subgroups, with the highest proportion of the groups being categorized as immune-ignored tumours. We further validated our approach using manual scoring based on the methodology of Hammerl et al. (2020) and found strong concordance with QuMix classifications, supported by significant agreement in concordance analysis (Figure 3d). To evaluate the prognostic significance of this classification, we fitted a Cox proportional hazards model including hormone receptor (HR) status, HER2 status, tumour grade, tumour stage, and tumour size as covariates. We found that QuMix classification identified immune-inflamed tumours were statistically associated with better prognosis compared to immune-ignored while immune-excluded had a lower hazard ratio but did not reach statistical significance (Figure 3e). In contrast, classification using the Hammerl method showed differences in hazard ratios that were not statistically significant.

### Machine learning–based integration of immune markers, clinical, and transcriptomic features for prognostic classification

To further assess the prognostic relevance of the QuPath-derived spatial features, we next tested if these features can stratify patients according to their prognosis. Using a 10-year cut-off, patients were first classified as having a good prognosis if they survived beyond 10 years, or a poor prognosis if they died within this period. We then used these features to train an ensembled machine learning model (described in our previous study) to classify patients into these prognostic groups. In brief, we adopted the same framework, comprising five iterations of stratified random train–test splitting (7:3) of the 145 patients (76 good prognosis, 69 poor prognosis) to generate independent training and testing datasets. With regard to predictive performance, the models trained using QuPath-derived features (CD8⁺ T-cell density at the invasive front and tumour center) demonstrated improved discriminative ability when combined with immunohistochemistry Aperio ImageScope-derived scores (CD3⁺, CD4⁺, and CD8⁺ T cells; CD3⁺/CD4⁺, CD3⁺/CD8⁺, and CD4⁺/CD8⁺ ratios; and PD-L1 expression), compared to when used alone (median AUROC = 0.71 [95% CI: 0.48–0.71] vs 0.59 [95% CI: 0.42–0.59]) (Table 1). In comparison, when spatial features derived from the Hammerl method were used instead, discriminative performance decreased (median AUROC = 0.54, 95% CI: 0.36–0.54), suggesting that QuPath-derived features are more prognostic (Table 1).This difference may reflect the underlying distinction in quantifying abundance, where the Hammerl method quantifies CD8⁺ infiltration as a percentage within predefined regions, whereas QuPath-derived features capture continuous cell density and spatial distribution across the tumour microenvironment. Therefore, the combination of QuPath-derived features and immunohistochemistry-derived scores (referred to in this study as digital pathology-derived immune features) was used in subsequent analyses.

**Table 1.** Development and evaluation of digital pathology-derived immune feature sets for prognostic classification. Performance of machine learning models trained using spatial features derived from QuPath, IHC-based immune markers, and their combinations. The area under receiver operating characteristic (AUROC) plot is reported as median with 95% confidence intervals.

| Feature Set | Description | AUROC (95% CI) |
| --- | --- | --- |
| **QuPath** | CD8<sup>+</sup> T-cell density at the invasive front and tumour center (delineate by QuPath) | 0.59 (0.42–0.59) |
| **IHC only** | CD3<sup>+</sup>, CD4<sup>+</sup>, and CD8<sup>+</sup> scores; CD3<sup>+</sup>/CD4<sup>+</sup>, CD3<sup>+</sup>/CD8<sup>+</sup>, and CD4<sup>+</sup>/CD8<sup>+</sup> ratios; and PD-L1 expression | 0.63 (0.43–0.63) |
| **QuPath + IHC (DP)** | Combination of QuPath and IHC | **0.71 (0.48–0.71)** |
| **Hammerl + IHC** | Percentage of CD8<sup>+</sup> T-cell at the border and center; ratio of the percentage at the border and center | 0.54 (0.36–0.54) |

To evaluate the prognostic value of digital pathology-derived immune features relative to established predictors, we compared their performance against models trained using clinical variables and a previously derived gene expression signature. The clinical variables were selected based on those incorporated in the PREDICT v3.0 model, excluding treatment and smoking status. The gene signature consisted of 77 genes identified through LASSO regression from transcriptomic profiles of 258 patients in our cohort, which includes all 145 patients included in this analysis.

Models trained using clinical variables alone achieved a median AUROC = 0.62, 95% CI: 0.37–0.62, Table 2). On the other hand, models trained and evaluated on the same subset of 145 patients using digital pathology-derived immune features alone achieved a median AUROC of 0.71 (95% CI: 0.48–0.71), outperforming models based on clinical variables. Notably, integrating digital pathology-derived immune features with clinical variables improved model performance from 0.62 to 0.68 (95% CI: 0.46–0.68), corresponding to an absolute gain of 0.07. This finding suggests that immune features derived from immunohistochemistry and spatial analysis provide prognostic information beyond conventional clinicopathological parameters.

**Table 2.** Comparative performance of digital pathology, clinical, and gene expression feature sets for prognostic classification. Performance of machine learning models trained using digital pathology-derived immune features (DP), clinical, and gene expression (GE) features, individually and in combination. The area under receiver operating characteristic (AUROC) plot is reported as median with 95% confidence intervals.

| Feature Set | Description | AUROC (95% CI) |
| --- | --- | --- |
| **Digital pathology (DP)** | Combination of QuPath and IHC features | 0.71 (0.48–0.71) |
| **Clinical** | Age at diagnosis, hormone receptor (ER/PR) status, HER2 status, tumour grade, tumour size, and number of locoregional lymph nodes to which cancer has spread | 0.62 (0.37–0.62) |
| **Gene expression (GE)** | 77-gene signature derived from 258 patients in the MyBrCa cohort, described in Khairi et. al. (2026) | 0.88 (0.75–0.88) |
| **DP + Clin** | Combination of DP and Clin | 0.68 (0.46–0.68) |
| **DP + GE** | Combination of QuPath and GE | 0.91 (0.79–0.91) |
| **Clin + GE** | Combination of Clin and GE | 0.87 (0.73–0.88) |
| **DP + Clin + GE** | Combination of all three features | 0.91 (0.80–0.92) |

Models trained using gene expression signatures demonstrated the strongest predictive performance, achieving a median AUROC of 0.88 (95% CI: 0.75–0.88). Addition of gene expression signatures to digital pathology-derived immune features only resulted in a modest improvement in discrimination, increasing the median AUROC to 0.91 (95% CI: 0.79–0.91), suggesting that digital pathology-derived immune features provide limited additional prognostic information beyond gene expression signatures in the same dataset.

Finally, models incorporating all three feature types: clinical, digital pathology-derived immune features, and gene expression, achieved a median AUROC of 0.91 (95% CI: 0.80–0.92). Collectively, these results demonstrate that immune-related digital pathology features enhance prognostication compared with clinical variables alone and provide complementary information to transcriptomic signatures, although incremental gain over gene expression-based models was modest.

## Discussion

In this study, we evaluated the prognostic utility of immune cell immunohistochemistry markers and spatial immune architecture in a large cohort of Asian breast cancer patients. Using semi-automated scoring of digitized whole slide images of breast tumour tissue samples stained for CD3, CD4, CD8, and PD-L1 markers, we observed that individual immune marker scores were not significantly associated with overall survival in 547 tumour samples. In contrast, when the spatial distribution of CD8⁺ cells was leveraged to classify immunophenotypes, tumours with the immune-inflamed phenotype were significantly associated with improved overall survival. We then assessed the discriminative performance of spatial metrics and immunohistochemistry scores within an ensemble machine learning framework and found that digital pathology–derived features achieved moderate performance in predicting 10-year prognosis.

### Biological and clinical relevance of immune IHC scores in our cohort

The positive correlation between CD3, CD4, and CD8 scores and overall survival has been seen in previous studies (Ali et al., 2014; Liu et al., 2011; Mahmoud et al., 2011; K. Wang et al., 2017). However, in this study, statistical significance was observed only among patients with the aggressive TNBC subtype, in whom higher CD4⁺ and CD8⁺ levels were associated with improved overall survival. This may be because TNBC is known to be more immunogenic than the other subtypes (Cardoso et al., 2019; Pan et al., 2020), and also suggests that immune activity is less relevant to patient outcomes in the breast cancer subtypes that are driven by hormonal and growth factors. High levels of these immune markers likely reflect a robust anti-tumour immune response and cytotoxic CD8⁺ T-cell–mediated tumour cell eradication (Raskov et al., 2021). These findings are in line with clinical trials for immunotherapy, where TNBC appears to have better response to immune checkpoint inhibitor therapy compared to other subtypes (Cardoso et al., 2019; Schmid et al., 2020).

In contrast, we did not find any association between PD-L1 and survival in the overall cohort, or among patients with TNBC. In contrast, previous meta-analyses of the prognostic significance of PD-L1 expression in breast cancer found that PD-L1 expression is associated with worse overall survival, although the 95% confidence intervals for the hazard ratios (HR) in these studies are close to or overlap with HR=1 (C. Wang et al., 2017). This discrepancy may be due to known population-specific differences in the tumour immune microenvironment (Pan et al., 2020), or differences in antibody or methodology. Of note, many of the individual studies included in the meta-analyses reported a lack of association with overall survival, suggesting that our study is within the range of previously reported results. Additionally, prior studies have often used more specific endpoints such as disease-free survival (DFS), breast cancer–specific survival, or treatment response measures including pathological complete response (pCR). This may partly explain the differences observed, as TILs are particularly associated with chemosensitivity and immune-mediated tumour control, especially in aggressive subtypes such as TNBC.

Similar to previous studies (Karn et al., 2017; McGrail et al., 2021; Safonov et al., 2017), we did not find any association between the immune markers with SNV and indel counts, TMB, or neoantigen load. Instead, we found that CD3, CD4 and CD8 scores were negatively correlated with CNAs across all breast cancer subtypes, albeit more strongly in TNBC, and across multiple CNA categories including loss, deletion, gain, and amplification. These results are in line with previous findings (Davoli et al., 2017; Karn et al., 2017; Lu et al., 2019; Zerdes et al., 2021) that found a negative correlation between somatic copy number alterations (SCNAs) and immune gene signatures in breast cancer. This negative correlation is likely due to a combination of focal SCNAs that drive cancer proliferation and immune evasion, pro-tumourigenic gene dosage effects from arm- and chromosomal-level SCNAs, as well as copy number loss and deletions in antigen presentation related genes (Davoli et al., 2017; Zerdes et al., 2021). In contrast, we found that CD4 scores were weakly positively correlated with HRD overall, whereas CD8 scores were negatively correlated with HRD in TNBC. Although HRD is often associated with higher mutational burden and neoantigen load (Budczies et al., 2022), our findings are consistent with previous research showing a negative correlation between HRD and immune scores in patients with germline BRCA mutations associated with NFκB activity (Kraya et al., 2019). Additionally, we found that the clock-like single-base-substitution (SBS) signatures 1 and 5 were negatively correlated with CD3 scores, while the APOBEC-associated SBS signatures 2 and 13 were positively associated with CD3 and CD4. The negative correlation with clock-like signatures is probably related to the decline of immune function with age (Foster et al., 2011), while the positive association with APOBEC-associated mutational signatures has been noted previously (Cescon et al., 2015; Wen et al., 2016) and may reflect APOBEC-driven somatic hypermutation (Nik-Zainal et al., 2014; Pan et al., 2021). Together, these results recapitulate many of the known associations between T-cell markers and prognosis in Asian breast cancer, and further support previous findings that these associations are largely restricted to the TNBC subtype.

In our data, all three immune markers had strong positive correlations with the cGAS-STING signalling, antigen processing and presentation, and T-cell receptor signalling pathways. These correlations were stronger in TNBC and weaker in the HR+ subtypes, which may be because TNBC is more genomically unstable (Chen et al., 2024; Curtis et al., 2012). The correlations with the antigen processing and presentation and T-cell receptor signalling pathways are expected as these are the canonical pathways by which T-cells are recruited into the tumour microenvironment, and represent additional confirmation that our results are robust. The association with cGAS-STING signalling is interesting as this suggests that the T cells in our cohort may be partially driven by the presence of cytosolic DNA (Kwon & Bakhoum, 2020). The recruitment of cytotoxic T cells by cGAS-STING signalling is well-established (Jiang et al., 2020), including in breast cancer (Chen et al., 2024); however, as T cells also express STING (Kuhl et al., 2023), further work will be required to establish the importance of cytosolic DNA in driving immune response in our cohort.

### Spatial immune profiling and its prognostic implications

Although the IHC-based CD8 marker did not achieve prognostic significance when analysed across all patients, classification of a subset of patients based on the spatial distribution of CD8⁺ cells revealed that the inflamed phenotype showed a significant association with overall survival. This is not unexpected, given different spatial metrics of CD8⁺ cells have been reported to be prognostic for different clinical outcomes (Niu et al., 2025). This underscores the importance of spatial information, as direct interactions between T cells and tumour cells are critical for cytotoxic activity and may not be captured by conventional IHC scoring. Additionally, given that H&E slides are routinely generated in clinical practice, these findings suggest that spatial analysis based on a single immune marker may still provide meaningful prognostic information.

Previous studies have applied the finite mixture modelling in cancer image analysis to investigate tumour heterogeneity by modelling histopathological image-environment interactions (Im et al., 2022; Im et al., 2023). While traditionally this analysis is used for clustering low-dimensional data such as the clinical, demographic and environmental data, or for risk stratification (Basu & Tiwari, 2010; Jia et al., 2022), more recent studies demonstrated its use in high-dimensional data such as RNA-seq (Prabakaran et al., 2019) and miRNA data (Hong et al., 2020) to predict recurrence. In this study, through the probabilistic finite mixture model-based classification of immunophenotype, we found that the largest proportion of patients in our dataset was classified as immune-ignored, even across different disease subtypes. This finding is consistent with the general understanding that breast cancer is a relatively immunologically “cold” tumour. Nevertheless, we observed a comparatively high proportion of inflamed tumours among patients with HR+/HER2− (32.7%) and HER2+ (40.0%) subtypes, which are generally considered to exhibit limited immune cell infiltration. In comparison, a relatively lower prevalence of immune-hot tumours has been reported, with approximately 7.8% in HR+/HER2− tumours in the METABRIC cohort (Griguolo et al., 2025), and 12.6% in HR+/HER2− tumours and around 19.0% in HER2+ tumours in the German cohort (Denkert et al., 2018). Our finding here is consistent with our previous study on a larger subset of this cohort, which revealed higher immune infiltration compared to Caucasian counterparts, characterized by elevated expression of immune-related gene signatures (Pan et al., 2020). Additionally, in another study, we found that the HR+/HER2− disease clusters based on immune profiles instead of the conventional luminal A-luminal B paradigm previously reported in breast cancers from women of European descent, further suggesting that immune activation may play a more important role in Asian HR+/HER2− breast cancer (Pan et al., 2024). Taken together, these findings highlight elevated immune infiltration and a distinct tumour–immune landscape in this population.

The QuMix pipeline demonstrated strong concordance with the manual Hammerl method in classifying immune-inflamed and immune-ignored phenotypes, with only partial concordance observed for the immune-excluded phenotype. Discrepancies across all phenotype classifications are likely attributable to the differences in spatial definitions between the Hammerl method and the QuMix pipeline, particularly in delineating tumour borders, invasive margins, and the tumour center. This, in combination with the whole-slide approach implemented in QuPath, may enable the capture of spatial features that are more strongly associated with prognosis compared to the Hammerl method, which relies on predefined spatial criteria that may be subject to inter-observer variability. This is further supported by the superior discriminative performance of machine learning models integrating QuPath-derived features compared to those using Hammerl-derived features, albeit in a smaller subset of patients. Taken together, these results support the validity of the QuMix pipeline and highlight the prognostic relevance of spatial cytotoxic TIL features in our cohort.

### Integration of immune and spatial features for prognostic assessment

In this study, an ensemble machine learning framework was used to integrate and evaluate the prognostic contribution of multiple immune markers and spatial features. Models trained on immunohistochemistry-derived scores showed moderate performance in stratifying patient survival at 10 years. Although individual TIL markers were not significantly associated with overall survival in adjusted Cox proportional hazards models, their integration in a multivariable model together with spatial information likely captures a more comprehensive immune profile, resulting in improved prognostic value. Interestingly, models trained on digital pathology-derived immune features outperformed those based on clinical variables, with only marginal improvement observed when the two were combined. This observation is consistent with studies in other cancer types (Lecuelle et al., 2024) and is not unexpected, as digital pathology can capture cell-level and spatial features that are more directly linked to tumour biology and prognosis. The performance of models trained on the gene expression signature is consistent with our previous study, which is expected as the current cohort includes a subset of patients from that dataset. Notably, integrating all three feature types did not further improve performance, suggesting that these models may have reached a performance plateau, as reflected by the unchanged AUROC. Together, these findings suggest that while gene expression contributes most strongly to discriminative performance, immune marker–derived features captured through digital pathology can still provide complementary information within a multimodal, integrative framework.

### Study limitations and conclusion

Several limitations of this study should be acknowledged. First, the MyBrCa cohort on which this study is based on was collected from participating private hospitals in urban areas of Malaysia and comprises a high proportion of patients of Chinese ethnicity. While this provides valuable insight into the tumour–immune landscape of underrepresented Asian breast cancer, it may limit the generalizability of our findings to the broader multiethnic Malaysian and Asian population. Another limitation relates to the use of finite mixture modelling to derive internal thresholds for immunophenotype classification. This method assumes multiple underlying subpopulations within the dataset. However, the unimodal nature of our data, indicating limited heterogeneity, may impact the robustness and interpretability of the resulting clusters. Next, the Hammerl method is inherently subjective, as it relies on qualitative definitions of tumour regions and immune cell distribution. While we followed the instructions described in the original Hammerl manuscript as closely as possible, the approach remains susceptible to inter-observer variability and our Hammerl quantifications should not be regarded as definitive. Finally, while our assessment of prognostic value for immune markers and subsequent spatial characterization of CD8⁺ cells was performed on a relatively large number of samples, the subsequent machine learning analysis was conducted on a smaller sample size due to data availability. This may increase the risk of overfitting and compromise model robustness, underscoring the need for validation in a larger and more diverse cohort.

In conclusion, our study demonstrates that TIL scores and PD-L1 expression achieve prognostic significance in Asian breast cancer patients when evaluated within a broader multimodal and spatial framework, rather than as individual markers. While prospective validation is warranted, these findings support the potential clinical utility of immune markers within a digital pathology framework that may be especially relevant in Asian breast cancer, which may have a more active immune landscape compared to their Western counterparts.

## Author Contributions

M.H.F.K. and J.-W.P. co-led the study design and data analysis. M.H.F.K. drafted the manuscript. H.M. contributed to data analysis. J.Y.T., P.-S.N., and C.W.K. contributed to data analysis and slide digitisation. S.N.H. contributed to sample collection and processing. I.Y.L., M.K., A.A., H.X.Y.L., C.-H.Y., and P.R. contributed to data acquisition and interpretation. S.-H.T. and J.-W.P. supervised the study. All authors reviewed, revised, and approved the final version of the manuscript.

## Acknowledgements

Genomic analyses were performed previously by the Genomics Core Facility at the Cancer Research UK Cambridge Institute, established in collaboration with Prof. Carlos Caldas, Dr. Chin Suet Feung, and Dr. Oscar Rueda. The authors gratefully acknowledge all patients who participated in the Malaysian Breast Cancer (MyBrCa) study and the clinical and laboratory teams at Cancer Research Malaysia and University Malaya Medical Centre for their invaluable contributions.

## Ethics declaration

Patient recruitment and sample collection was reviewed and approved by the Independent Ethics Committee, Ramsay Sime Darby Health Care (Reference nos: 201109.4 and 201208.1), as well as the Medical Ethics Committee of the University Malaya Medical Centre (Reference no: 842.9). Written informed consent to participation in research was given by each individual patient. All procedures involving human participants were conducted in accordance with the ethical standards of the institutional research committees and with the principles of the International Council for Harmonisation E6 Good Clinical Practice (ICH E6 GCP) guidelines.

## Data availability

The datasets supporting the conclusions of this article are available in the European Genome-phenome Archive repository [Accession no. <u>EGAD00001009311</u>]. Access to controlled patient data will require the approval of the Data Access Committee. Further information is available from the corresponding author upon request.

## Abbreviations

AUROC: Area Under the Receiver-Operator Characteristic curve
TILs: Tumour-infiltrating lymphocytes

