## Supplemental Figures S1-S5 for "Prognostic utility of spatial immunophenotyping in Asian breast cancer"

### Supplementary Figures

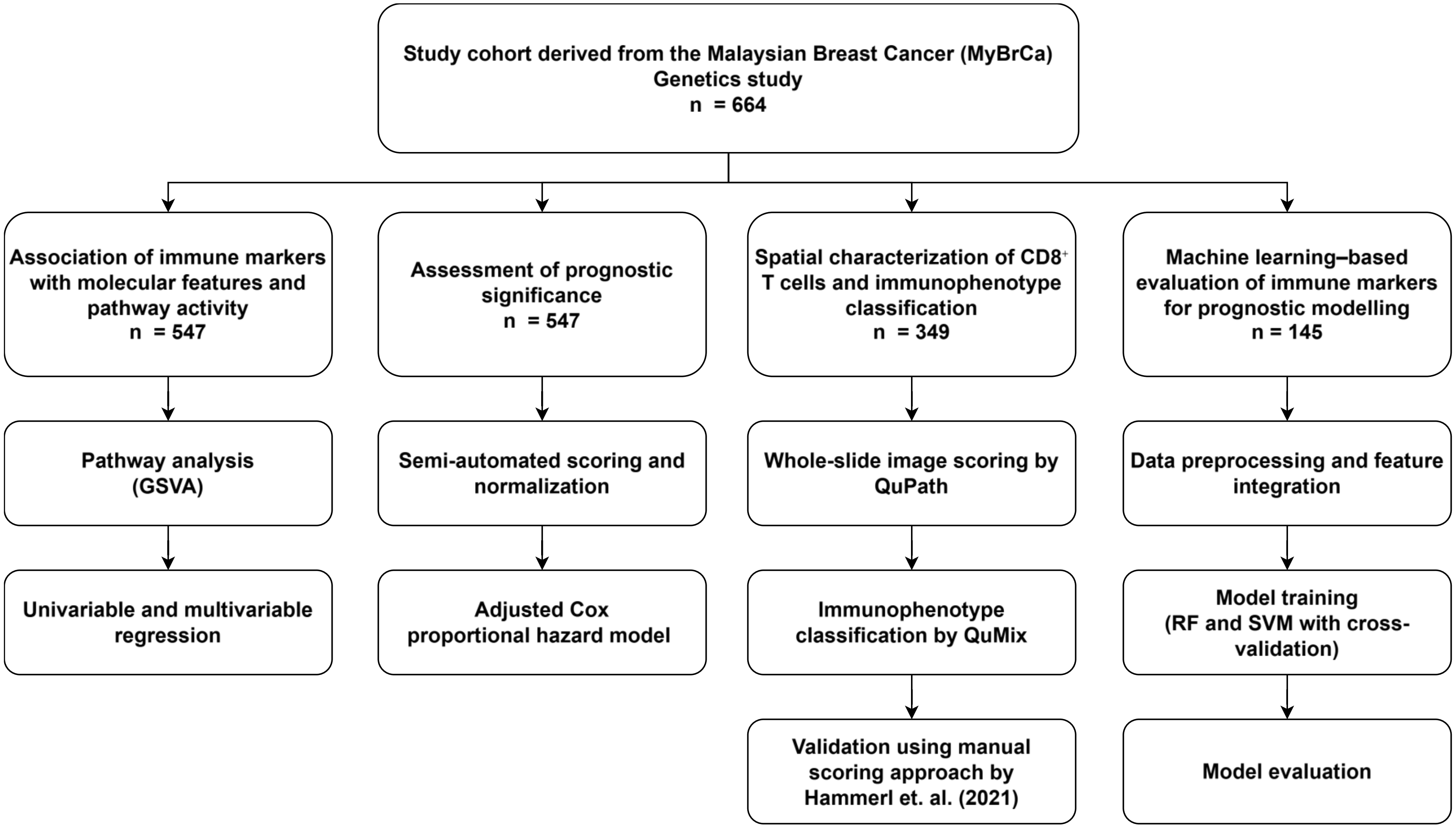


**Figure S1 Schematic overview of the study design and the analysis workflow**

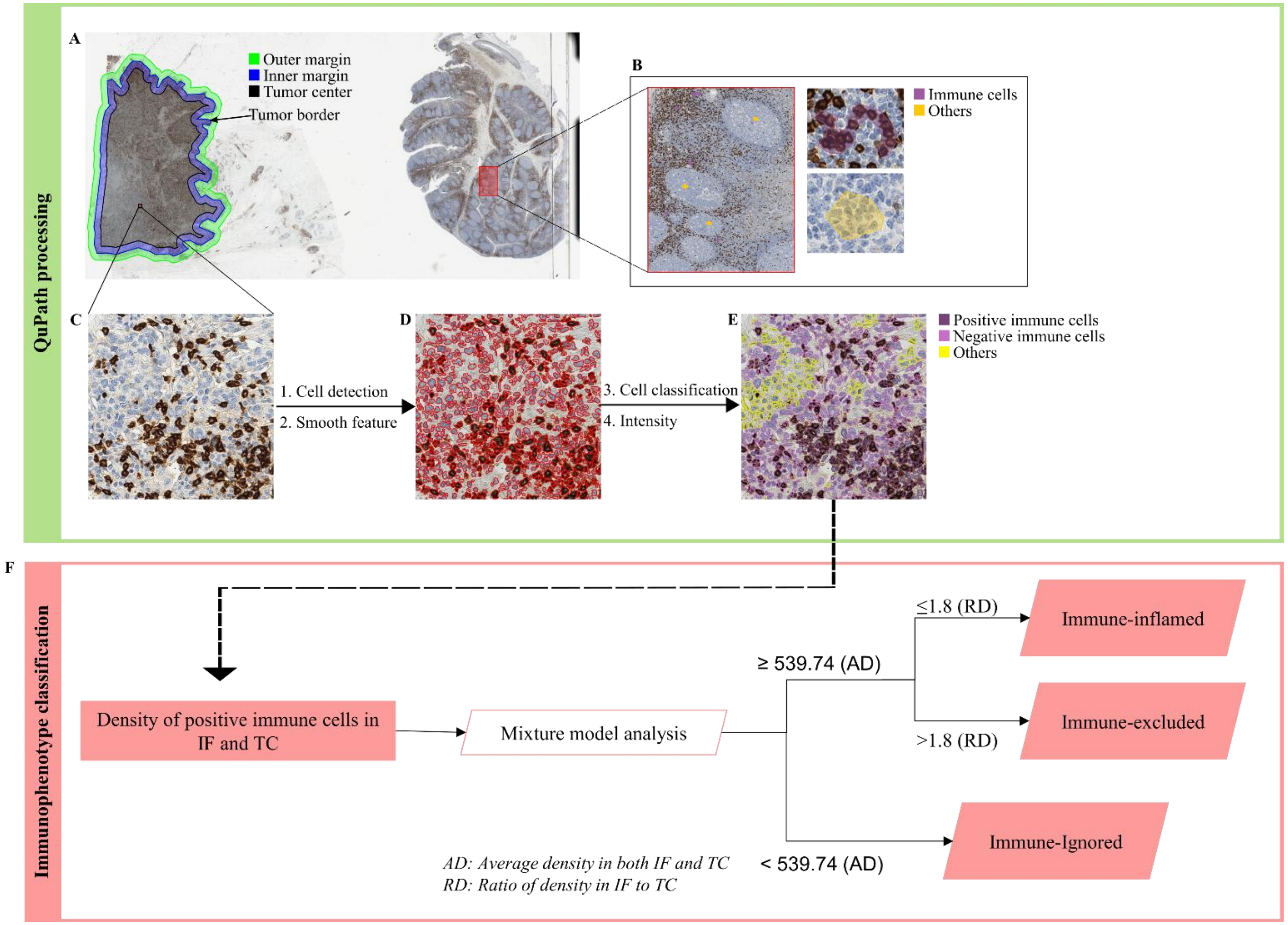


**Figure S2 Overview of QuMix pipeline in which QuPath processing integrated with mixture model analysis.** A, tumour border was manually created and then divided into three regions: outer margin, inner margin and tumour center (TC). Outer and inner margin were defined as invasive front (IF). B, a region within the control tissue used as a training for classification of immune cells and others. Example of QuPath processing flow from before cell segmentation (C), to segmented cells in red color (D) and finally with cells classified into positively- and negatively-stained immune cells and other cells (E). F, immunophenotypes classification into immune-ignored, immune-excluded and immune-inflamed using thresholds generated by two-component mixture model.

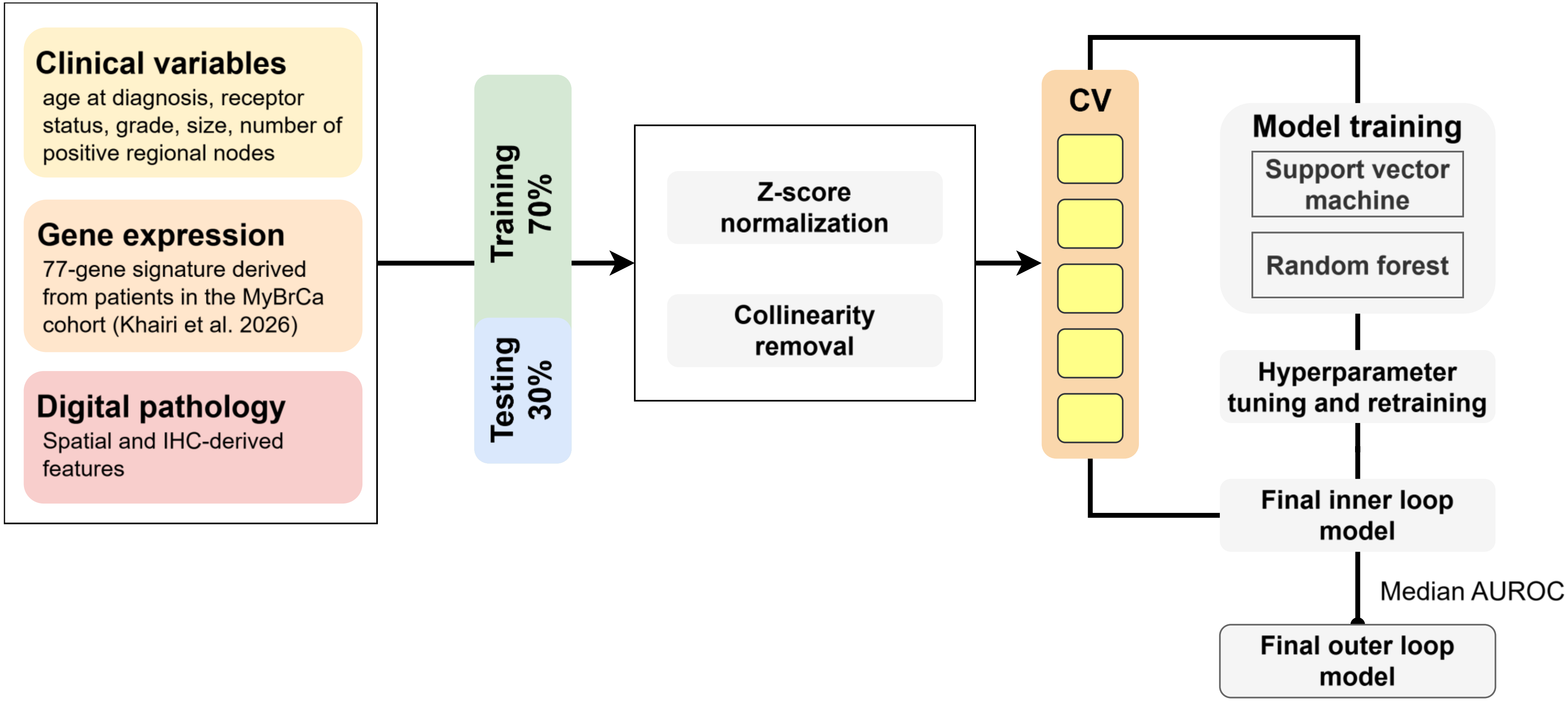


**Figure S3 Schematic overview of the ensemble machine learning framework.** Adapted from Khairi et. al (2026). Clinical, gene expression, and digital pathology-derived immune features were split into training (70%) and testing (30%) sets. Training data were standardized and filtered for collinearity, followed by five-fold cross-validation for model training, hyperparameter tuning, and retraining using Random Forest and Support Vector Machine algorithms. Predictions were aggregated to form a final inner loop model for each split. Across five iterations, the model with the median AUROC on the test sets was selected as the final outer loop model. AUROC, area under the receiver operating characteristic curve; CV, cross-validation.

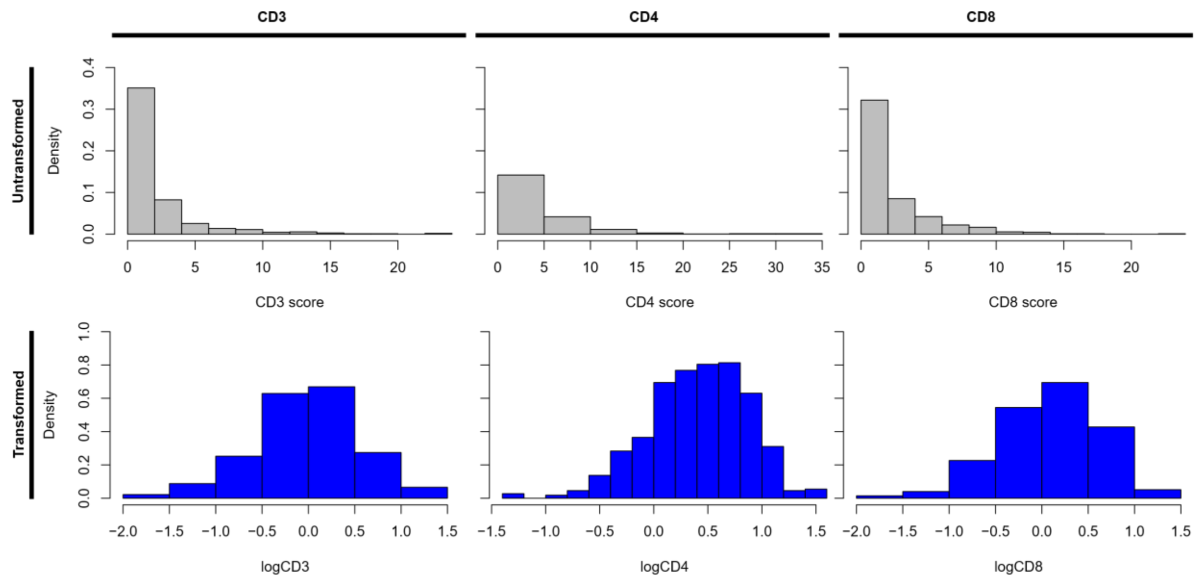


**Figure S4 Distribution of immune marker scores (CD3, CD4, and CD8) before and after log transformation.** Distributions are shown for both untransformed and log-transformed values to illustrate normalization towards a Gaussian-like distribution.

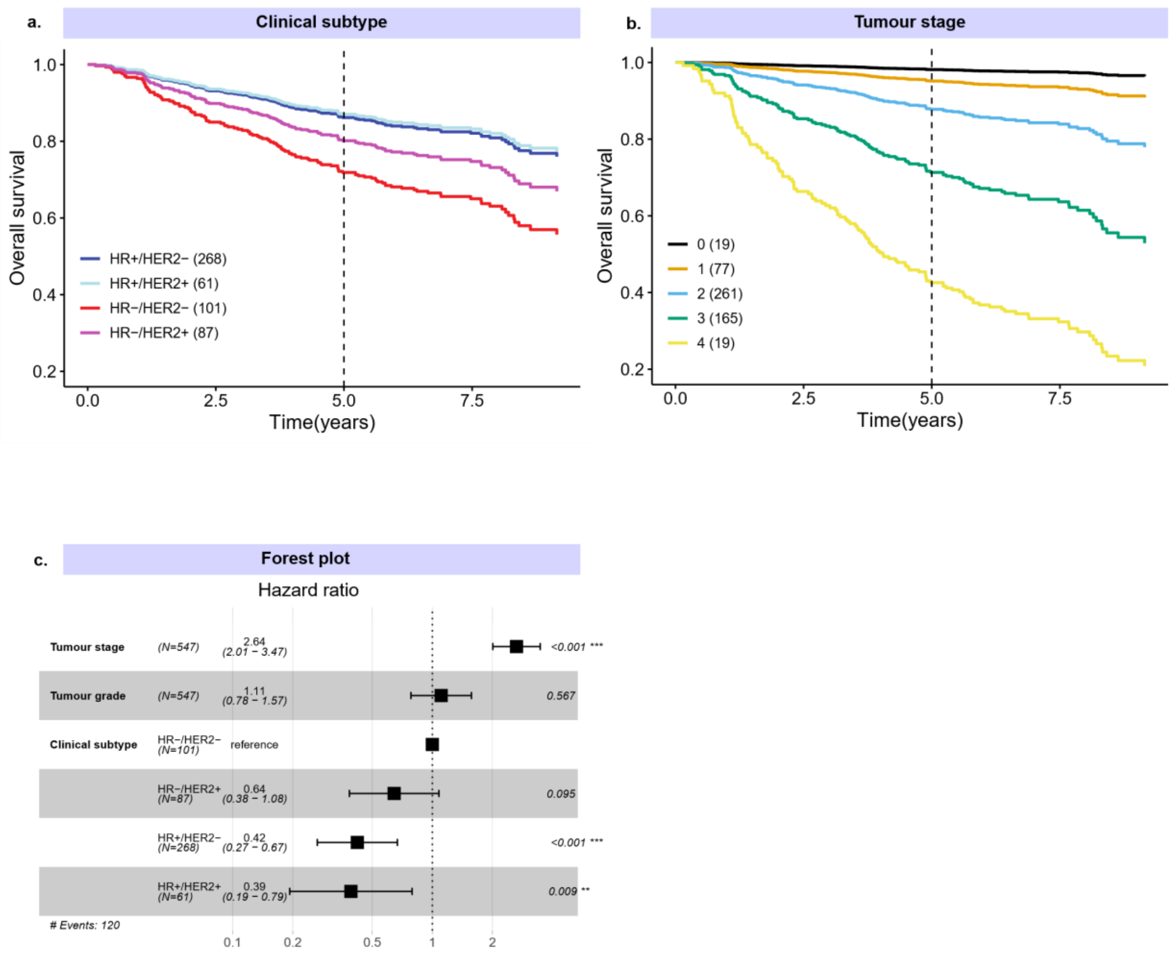


**Figure S5 Kaplan–Meier survival analysis and hazard ratios across clinical subtypes and tumour stages.** a–b) Kaplan–Meier curves showing overall survival stratified by clinical subtype and tumour stage. c) Forest plot of adjusted hazard ratios (HRs) for overall survival derived from a multivariable Cox proportional hazards model. The model includes tumour stage, tumour grade, and clinical subtype as covariates.
